# Glucocorticoid Receptor ablation promotes cardiac regeneration by hampering cardiomyocyte terminal differentiation

**DOI:** 10.1101/2020.01.15.901249

**Authors:** Nicola Pianca, Francesca Pontis, Maila Chirivì, Valentina Papa, Luca Braga, Rahul Shastry Patnala, Chiara Bongiovanni, Martina Mazzeschi, Kfir-Baruch Umansky, Giovanna Cenacchi, Mattia Lauriola, Mauro Giacca, Roberto Rizzi, Eldad Tzahor, Gabriele D’Uva

## Abstract

In mammals, glucocorticoid levels rise dramatically shortly before birth and prepare the foetus for post-natal life by promoting the maturation of the lungs and other organs. However, their impact on cardiac postnatal growth and regenerative plasticity is unknown.

Here, we demonstrate that exposure to endogenous glucocorticoids facilitates cell cycle exit and reduces the proliferation of neonatal cardiomyocytes. This cytostatic activity is shared by several synthetic glucocorticoid receptor (GR) agonists routinely used in clinical settings. We also observed that GR levels increase in cardiomyocytes during early post-natal development. Importantly, *in vivo* cardiomyocyte-specific GR ablation delayed the transition from hyperplastic (increase in cell number) to hypertrophic (increase in cell size) growth. Further, GR ablation partially impaired cardiomyocyte maturation, reducing myofibrils-mitochondria organization along with the expression of genes involved in fatty acid metabolism, mitochondrial respiration and energy transfer from mitochondria to the cytosol. Finally, we show increased cardiomyocyte proliferation in GR ablated juvenile and adult cardiomyocytes in response to myocardial infarction *in vivo*, thus promoting cardiac tissue regeneration.

We suggest that GR antagonization could serve as a strategy for heart regeneration based on endogenous cardiomyocyte renewal.

## INTRODUCTION

In mammals, severe cardiac injuries, such as those induced by myocardial infarction, result in a significant loss of cardiac muscle cells (cardiomyocytes), which are replaced by non-contractile fibrotic scar tissue. The loss of contractile cells and the inability of the adult mammalian heart to generate new cardiomyocytes is a major cause of cardiac dysfunction and heart failure ^1–4^.

It has been discovered that the neonatal mammalian heart is able to respond to an injury by a robust regenerative process ^5–7^. This regeneration is carried out by existing cardiomyocytes that undergo partial sarcomere disassembly and proliferate ^5^. Nevertheless, shortly after birth, most cardiomyocytes in the mammalian heart permanently exit from the cell cycle ^8,9^, and lose their ability to promote regeneration upon damages ^5^. The adult mammalian heart exhibits a very slow rate of cardiomyocyte renewal, clearly insufficient for a robust regenerative response ^10,11^. Improving the proliferative ability of endogenous adult cardiomyocytes represents an emerging strategy to repair the heart tissue and to boost cardiac function in heart failure patients ^2–4,12–14^.

Glucocorticoids are steroid hormones synthesised in the adrenal cortex and released into the circulatory system, exerting most of their actions through the Glucocorticoid Receptor (GR) ^15–17^. GR resides and gets activated directly in the intracellular space, then translocates to the nucleus and functions as ligand-activated transcription factors, hence the name “nuclear receptor”.

Physiological glucocorticoids (predominantly cortisol in mammals, including humans, and corticosterone in amphibians, reptiles, birds and rodents, including laboratory mice) influence many physiological processes, including development, immunity, inflammation, and stress response ^15,16^. However, surprisingly little is known about the direct role of glucocorticoid signalling in cardiac health and disease ^16,18,19^.

In mammals, glucocorticoids rise dramatically shortly before birth and are known to promote the maturation of the lungs and other organs to prepare the foetus for the post-natal life ^20,21^. Despite several studies suggesting the involvement of glucocorticoid signalling in foetal heart maturation in mice ^18,19,22,23^, the role of glucocorticoids in post-natal cardiomyocyte growth and regenerative plasticity is currently unknown.

By combining developmental-, cell- and molecular biology approaches, we analysed the impact of the glucocorticoid receptor in the regulation of postnatal cardiomyocyte maturation and growth, along with its impact on cardiac regenerative ability. We demonstrate that endogenous and synthetic glucocorticoids restrain the proliferative ability of immature neonatal cardiomyocytes. We also show that GR abundance increases in cardiomyocytes during early post-natal development. Importantly, cardiomyocyte-specific GR ablation (GR-cKO) delayed the postnatal transition from hyperplastic growth (cell proliferation) to hypertrophic growth (increase in cell size). RNA sequencing and gene ontology analysis coupled with ultrastructure examination by transmission electron microscopy, demonstrated that GR ablation impairs postnatal cardiomyocyte maturation, by reducing the expression of genes involved in fatty acid oxidation machinery, mitochondrial respiration capacity and energy transfer from mitochondria to the cytosol, as well as mitochondria-myofibrils organization. Finally, upon myocardial infarction GR ablated juvenile and adult cardiomyocytes were more proliferative, thus promoting cardiac tissue regeneration with reduced scar formation. Altogether these results support a model where a postnatal GR rise restrains the regenerative plasticity of cardiomyocytes. We thus propose that the modulation of GR activity in adult cardiomyocytes may represent a promising therapeutic approach for cardiomyocyte replacement upon cardiac injuries.

## RESULTS

### Endogenous and synthetic glucocorticoids restrain the proliferative ability of neonatal cardiomyocytes

We first analysed the effect of corticosterone, the main endogenous glucocorticoid in rodents, on the proliferation of neonatal cardiomyocytes (post-natal day 1, P1). Indeed, cardiomyocytes at this developmental stage retain a certain degree of proliferation, reminiscent of the embryonic stage. Cardiomyocyte proliferation was assessed by co-immunostaining with antibodies for cardiomyocytes, namely cardiac troponin T (Tnnt2) or cardiac troponin I (Tnni3), together with proliferative markers, namely Ki67 for cell-cycle re-entry and BrdU incorporation for DNA Synthesis (S) phase. *In vitro* administration of corticosterone promoted a remarkable dose-dependent decrease in cardiomyocyte proliferation, as shown by immunostaining for Ki67 and BrdU (**Fig. 1a,b**). Cardiomyocytes can undergo division of the nucleus (karyokinesis) without cytoplasm division (cytokinesis), which eventually leads to binucleation, a process linked to cardiomyocyte terminal differentiation. Thus, we also examined the effect of corticosterone on the final abscission of the two daughter cardiomyocytes by the immunostaining of Aurora B kinase, which localizes at the central spindle during anaphase and at the cleavage furrow during cytokinesis. Importantly, this analysis showed that the rate of cardiomyocyte cytokinesis was robustly inhibited by the administration of intermediate doses of corticosterone and completely suppressed by high doses (**Fig. 1c**).

**Figure 1.**
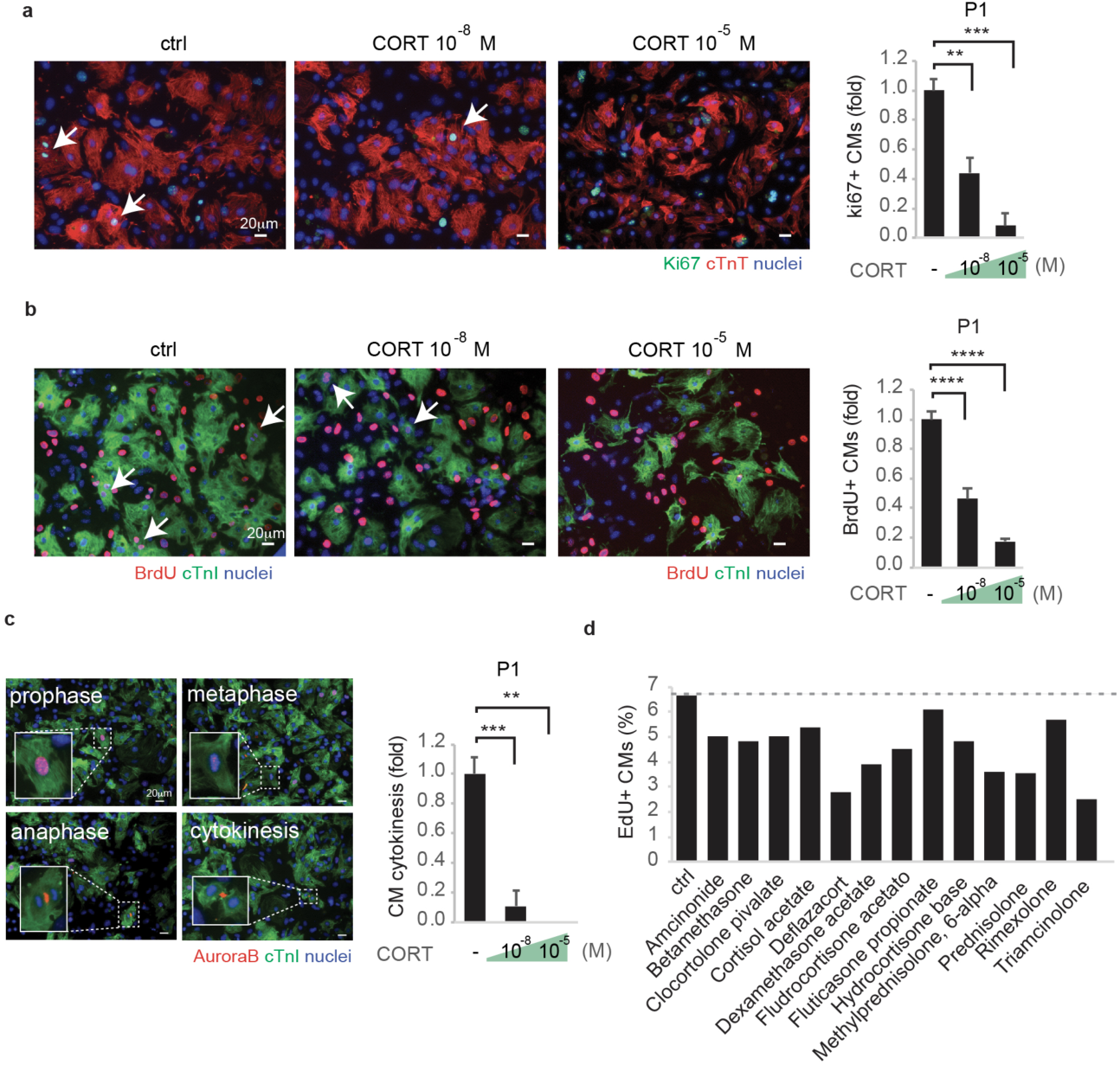
Exposure to endogenous and synthetic glucocorticoids induces cell cycle exit and reduces the proliferation of neonatal cardiomyocytes. **(a-c**) 1-day-old (P1) mouse cardiomyocytes (CMs) were cultured *in vitro* and stimulated with various doses of corticosterone (CORT); CMs were identified by cTnI or cTnT staining (see Methods for further details) and analysed by immunofluorescence for cell-cycle activity (Ki67, **a**), DNA synthesis (BrdU, **b**) and cytokinesis (Aurora B kinase, **c**); The proliferative rate of untreated control (ctrl) CMs was 3.6 +/- 0.9 %, 6.4 +/- 0.9 % and 0.34 +/- 0.1 % for ki67+, BrdU+ and mid-body Aurora B+ events, respectively; representative pictures are provided; arrows point at proliferating CMs; scale bars, 20 μm; (**d**) neonatal rat cardiomyocytes were cultured *in vitro*, stimulated with FDA approved GR agonists and analysed for DNA synthesis (EdU). In all panels, numerical data are presented as mean (error bars show s.e.m.); * p < 0.05, ** p < 0.01, *** p < 0.001 and **** p < 0.0001.

Synthetic glucocorticoids are widely used in the clinic as anti-inflammatory agents ^24,25^, however their direct effects on cardiomyocyte proliferative ability were poorly explored. Thus, we screened FDA-approved active agonists of the Glucocorticoid Receptor (GR) on cardiomyocyte cell cycle activity by EdU incorporation assay. Strikingly, our results show that all tested drugs reduced the proliferation of cardiomyocytes (**Fig. 1d**). These data suggest that endogenous glucocorticoids, as well as clinically relevant GR agonists, facilitate cell cycle exit and reduce the proliferation of immature cardiomyocytes in the early post-natal period.

### Cardiac GR abundance increases in cardiomyocytes during the early post-natal development

Nuclear localization of GR is associated with positive transcriptional activity. In order to clarify the activation status of this receptor *in vivo* during early post-natal development, we analysed GR nuclear immunoreactivity in heart sections from mice at P1 and P7. GR appeared localized in the nucleus in the majority of cardiomyocytes both at P1 and P7 stages (**Supplementary Fig. 1a**), suggesting that the receptor is active at both stages.

We next examined the dynamic regulation of GR expression levels within the cardiac tissue during the early post-natal period. To this end, we evaluated GR levels in heart lysates near birth (P0-P1) and one week later (P7). Intriguingly, our data revealed that cardiac GR expression increases during the first week of post-natal life, both at mRNA and protein levels (**Fig. 2a, b**). To analyse if the increase in GR abundance from P1 to P7 was attributable to a change in expression occurring in cardiomyocytes or in other cardiac stromal cells, cardiomyocytes were separated from cardiac stromal cells by immuno-magnetic cell sorting (see methods for further details) and gene expression analysis was performed. We first confirmed that the cardiomyocyte-specific marker Troponin T was more abundant in the cardiomyocyte fraction (**Supplementary Fig. 1b**), and that markers of other cardiac cell types, such as PECAM for endothelial cells and DDR2 for fibroblasts, were almost exclusively expressed in the stromal subset (**Supplementary Fig. 1c-d**), validating the specificity of the tissue fractionation procedure. Importantly, the analysis of GR expression levels demonstrated that the increase in GR expression during the early neonatal period occurs specifically in cardiomyocytes and not in other cardiac stromal cells (**Fig. 2c**). These data suggest that, in addition to the well-known increase in circulating glucocorticoids occurring in the immediate pre-birth period, an increase in the receptor abundance in cardiomyocytes likely boosts this signalling pathway in the early post-natal period (**Fig. 2d**).

**Figure 2.**
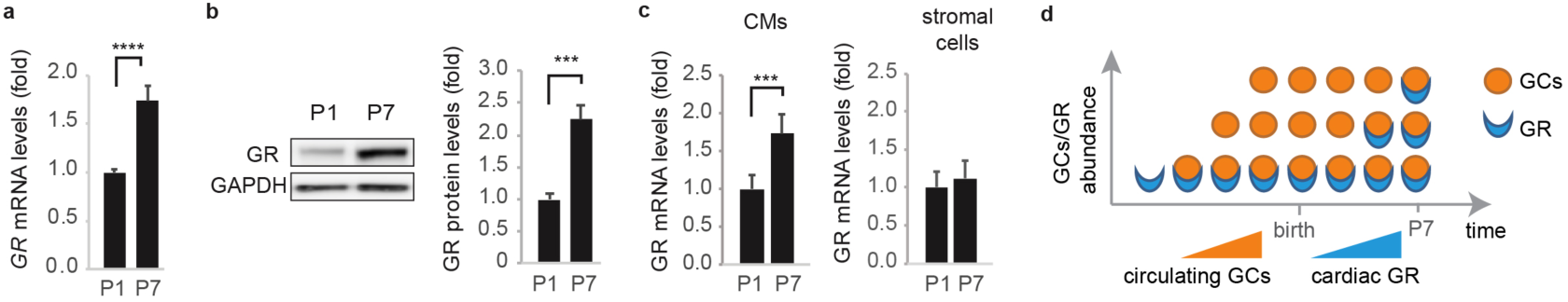
Glucocorticoid Receptor (GR) abundance increases in cardiomyocytes during the early post-natal period. **(a**) mRNA expression levels of Glucocorticoid Receptor (GR) in postnatal day 1 (P1) and postnatal day 7 (P7) heart lysates; (**b**) Western blot analysis of GR in P1 and P7 heart lysates; representative pictures and quantifications are provided; (**c)** GR mRNA expression levels in cardiomyocytes and stromal cells isolated from postnatal day 1 (P1) and postnatal day 7 (P7) hearts; (**d**) schematic diagram showing increased GCs/GR axis activation during early postnatal period, as result of the pre-natal increase in circulating glucocorticoids levels coupled with the post-natal increase in glucocorticoid receptor. In all panels, numerical data are presented as mean (error bars show s.e.m.); * p < 0.05, ** p < 0.01, *** p < 0.001 and **** p < 0.0001.

### Cardiomyocyte-specific deletion of Glucocorticoid Receptor (GR-cKO) increases the proliferation rate of neonatal cardiomyocytes

To evaluate the direct role of GR on cardiomyocyte proliferation, we generated a cardiomyocyte-specific GR conditional knock-out mouse model (GR-cKO mice). GR-cKO mice were previously generated and reported to appear normal till adulthood ^26^, however a potential impact on cardiomyocyte proliferation was not evaluated. We crossed floxed GR mice (GR-loxP/loxP), which contain loxP sites flanking exons 3 of the GR gene, with mice expressing Cre recombinase under the control of the cardiac alpha myosin heavy chain promoter (αMyh6-Cre +/-) (**Fig. 3a**). Cardiac GR ablation was confirmed at the protein level (**Fig. 3b**). Further, immunofluorescence analysis of cardiomyocytes isolated from neonatal GR-cKO mice confirmed that the deletion occurred specifically in cardiomyocytes (**Supplementary Fig. 2a**).

**Figure 3.**
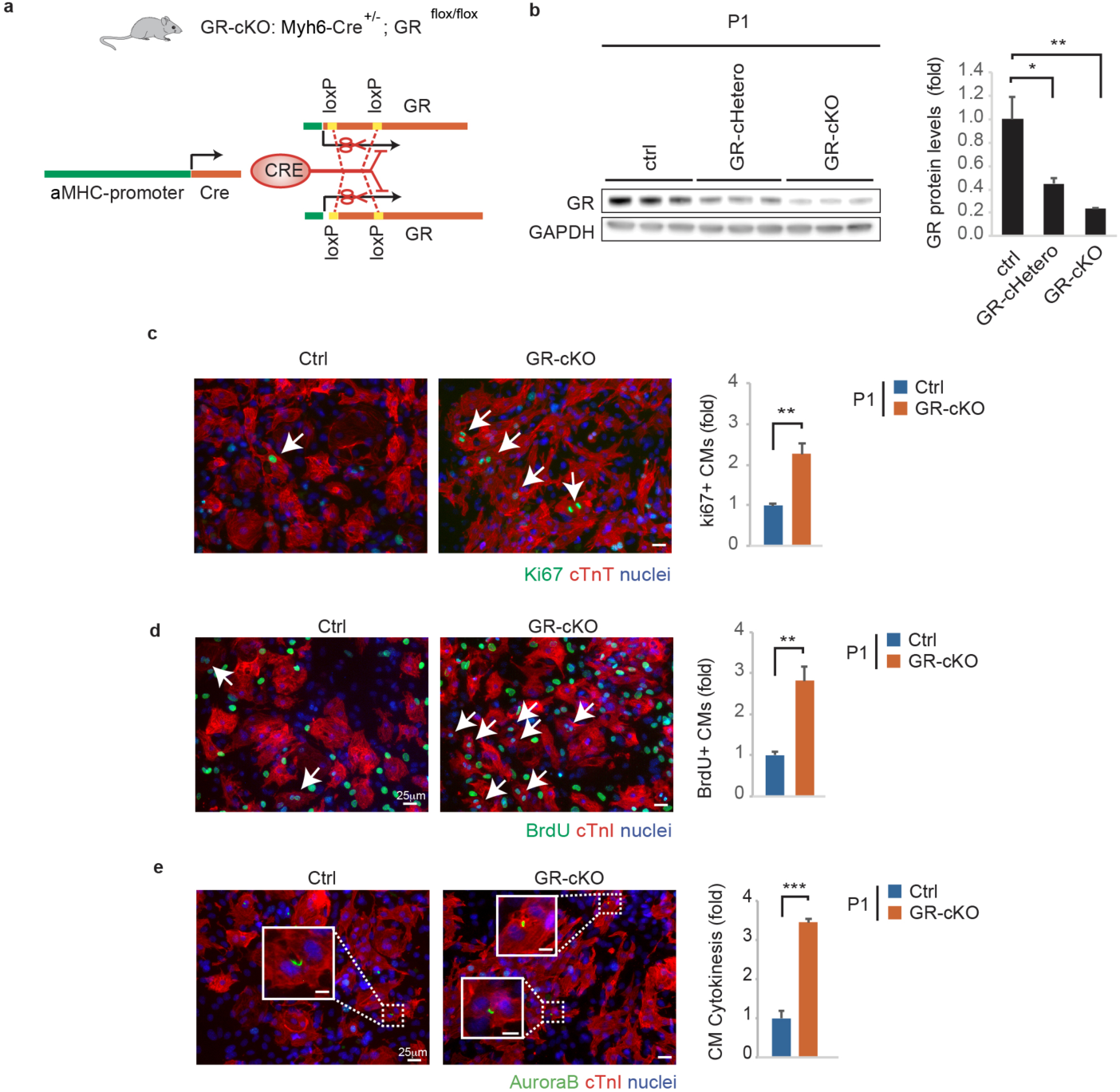
*In vitro* proliferation of neonatal cardiomyocytes is increased by cardiomyocyte-specific deletion of the Glucocorticoid Receptor (GR-cKO). **(a**) A schematic diagram depicting the generation of cardiomyocyte-restricted GR knock-out (GR-cKO) mice; (**b**) Western Blot analysis of GR protein levels in neonatal (P1) control (ctrl), GR-Hetero and GR-cKO heart lysates; (**c-e**) Immunofluorescence analysis of cell-cycle activity (Ki67, **c**), DNA synthesis (BrdU, **d**) and cytokinesis (Aurora B kinase, **e**) in cardiomyocytes isolated from P1 ctrl and GR-cKO mice; representative pictures are provided; arrows point at proliferating cardiomyocytes; scale bars, 25 μm. In all panels, numerical data are presented as mean (error bars show s.e.m.); * p < 0.05, ** p < 0.01, *** p < 0.001 and **** p < 0.0001.

Importantly, P1 GR-cKO cultured cardiomyocytes displayed higher proliferation rates compared to the control counterpart, in terms of cell cycle re-entry (**Fig. 3c**), DNA synthesis (BrdU incorporation, **Fig. 3d**) and cell division (cytokinesis, **Fig. 3e**). These data clearly demonstrate that GR has a key role in repressing the proliferative ability of immature neonatal cardiomyocytes.

### GR-cKO delays the early postnatal transition from hyperplastic to hypertrophic growth

During embryonic development in mammals, cardiomyocytes actively proliferate to build a functional heart, a phenomenon known as hyperplastic growth. During the early postnatal development, coincident with the first week of postnatal life in mice, most cardiomyocytes withdraw from the cell cycle and continue to grow in size (hypertrophic growth) ^8,9^. The evidence of the increase in GR abundance in cardiomyocytes during the early postnatal period (see **Fig. 2**), along with the robust inhibitory activity of GCs/GR axis on neonatal cardiomyocyte proliferation (see **Figs. 1-3**) prompted us to investigate whether GR could play a role in the physiological transition from hyperplastic to hypertrophic growth occurring shortly after birth.

As a first indication of cardiac growth upon cardiomyocyte-specific GR ablation, we measured the heart weight normalized to body weight (HW:BW ratio) of GR-cKO compared to control mice. No differences were appreciable at birth, whereas at postnatal-day-7 (P7) a small but statistically significant increase in HW:BW ratio was observed in GR-cKO (**Fig. 4a, Supplementary Fig. 2b**). Importantly, GR-cKO cardiomyocytes displayed reduced size (**Fig. 4b**). The slight increase in HW:BW ratio coupled with the reduction in cardiomyocyte size, suggests that cardiomyocyte number is increased in GR-cKO hearts. In line with this hypothesis, immunofluorescence analysis revealed an increased proliferation of cardiomyocytes in P7 GR-cKO hearts compared to controls (**Fig. 4c-d**).

**Figure 4.**
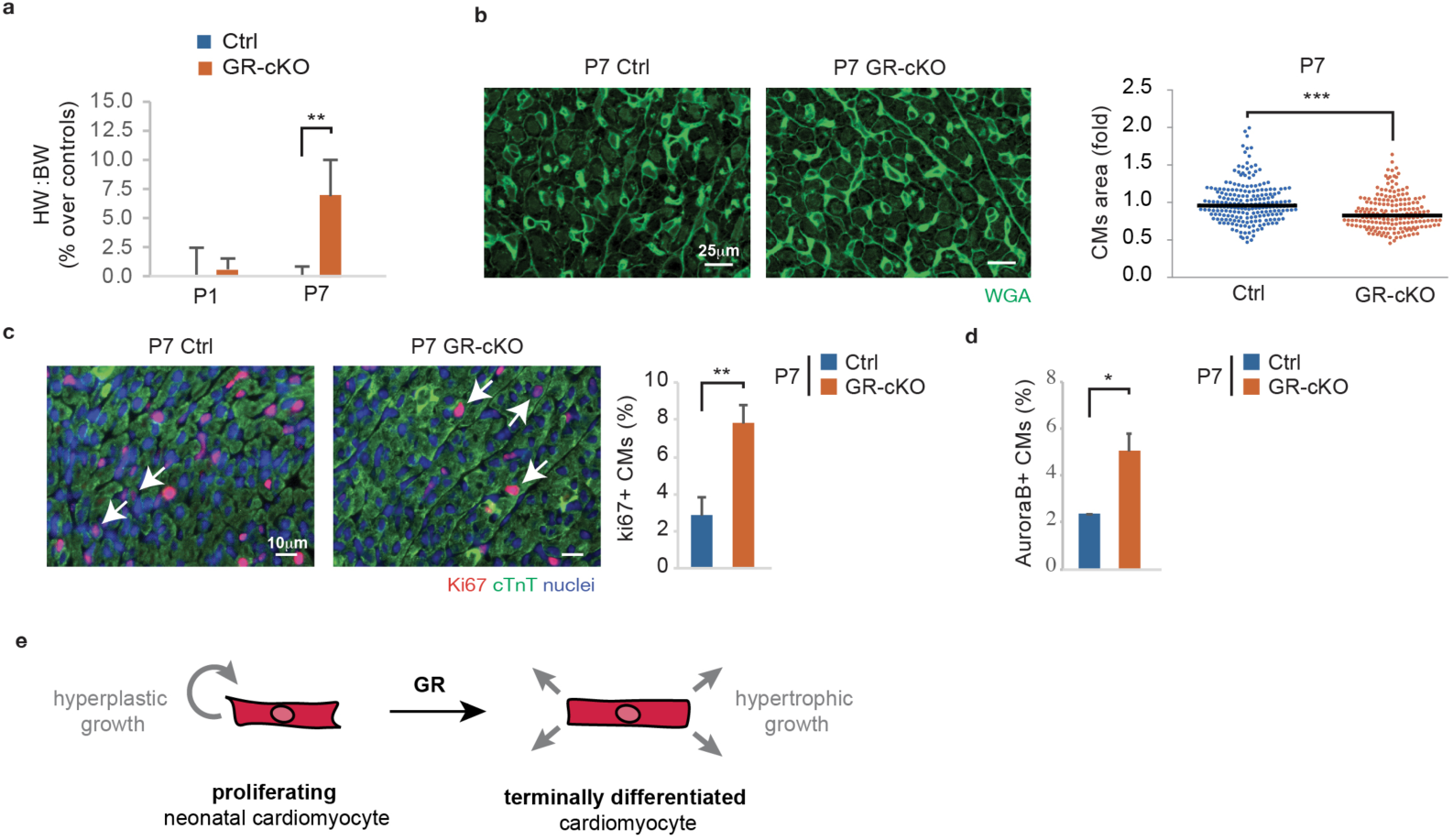
GR-cKO reduces the hyperplastic to hypertrophic transition occurring during the early postnatal period. **(a)** Percentage change of Heart Weight/Body Weight ratio (HW/BW) in control and GR-cKO animals at different stages of postnatal life (P1 and P7); (**b)** cardiomyocyte cross-sectional area evaluation by immunofluorescence analysis of wheat germ agglutinin (WGA) on P7 control and *GR*-cKO ventricular heart sections; representative images are provided; **(c-d**) *In vivo* evaluation of cardiomyocyte (CM) proliferation in control and GR-cKO ventricular heart sections; CMs were identified by cTnI or cTnT staining (see Methods for further details), proliferation was evaluated by immunofluorescence analysis of Ki67 (**c**) and Aurora B kinase (**d**). Representative images are provided, arrows point at proliferating CMs; (**e**) Schematic diagram showing the role of GR in promoting the postnatal developmental transition of neonatal cardiomyocyte from hyperplastic to hypertrophic growth. In all panels, numerical data are presented as mean (error bars show s.e.m.); * p < 0.05, ** p < 0.01, *** p < 0.001 and **** p < 0.0001.

These results show that the deletion of GR prolongs the post-natal proliferative window of cardiomyocytes while reducing the hypertrophic growth, suggesting an important role for GR in the transition from hyperplastic to hypertrophic growth (**Fig. 4e**).

### GR-cKO reduces the post-natal cardiomyocyte metabolic and sarcomere maturation

To shed light on molecular alterations driven by the GR ablation, we performed RNA-sequencing analysis on P7 GR-cKO and control heart lysates. As expected, GR was the most significant downregulated gene in GR-cKO hearts (**Supplementary Fig. 3**). Gene ontology analysis of these data, highlighted major changes in gene networks linked to metabolism and sarcomere organization.

Indeed, gene ontology analysis according to biological processes, pointed out that significant changes were observed in gene networks connected with cellular lipid metabolic and catabolic processes (**Fig. 5a**, heatmaps in **Supplementary Figure 4a-b**), in particular those related to fatty acid metabolic and biosynthetic processes (**Fig. 5a**, heatmap in **Supplementary Figure 4c-e**). Importantly, genes participating in fatty acid oxidation were all significantly downregulated in GR-cKO hearts (**Fig. 5b**). Consistent with the role of GR in lipid metabolism, gene ontology analysis according to molecular function highlighted a role in lipase activity (**Fig. 5c**, heatmap in **Supplementary Figure 5**). Furthermore, the expression of genes participating to electron transfer activities in mitochondria was also significantly reduced in GR-cKO hearts (**Fig. 5d**), including the Electron Transfer Flavoprotein (ETFA, which catalyses the initial step of the mitochondrial fatty acid beta-oxidation), Electron Transfer Flavoprotein Dehydrogenase (ETFDH, an inner mitochondrial membrane protein that passes electrons to ubiquinone), along with enzymes of respiratory Complex II (Succinate dehydrogenase complex subunit C - SDHC), Complex III (Ubiquinol-cytochrome c reductase - UQCRFS1), and Complex IV (cytochrome c oxidase - COX7B, COX7C), as well as cytochrome c (CYCS, which transfers electron between Complexes III and IV). Finally, we observed that mitochondrial creatine kinase (CKMT2), the enzyme that is responsible for the transfer of high energy phosphate from mitochondria to the cytosolic carrier creatine, was among the top 10 downregulated genes in GR-cKO hearts (see **Supplementary Fig. 3a)**.

**Figure 5.**
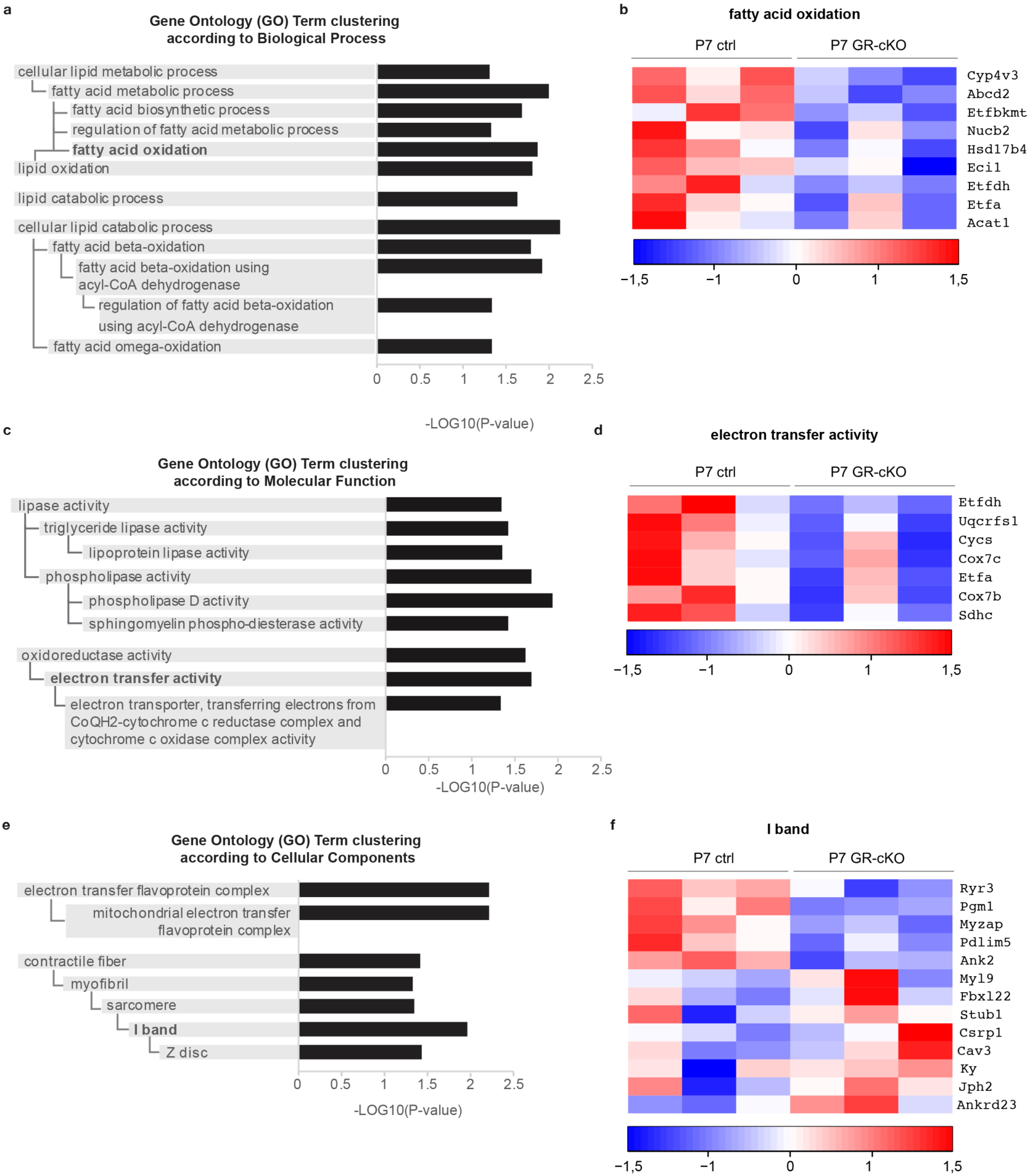
Gene ontology analysis highlights a role for GR in controlling genes involved in fatty acid metabolism, mitochondrial respiration and contractile fibers. **(a,c,e)** Gene ontology analysis according to (**a**) Biological Process, (**c**) Molecular Function and (**e**) Cellular Components of RNA-Seq data of P7 GR-cKO versus control hearts; (**b,d,f**) Heatmaps of differentially expressed genes by RNA-Seq transcriptome analysis of P7 GR-cKO versus controls hearts according to the following gene ontology terms: fatty acid oxidation (GO:0019395, **b**) electron transfer activity (GO:0009055, **d**) and I band (GO:0031674, **f**).

Beginning in late gestation and continuing through the first weeks after birth, cardiomyocytes undergo an important transition from anaerobic glycolysis in the cytosol to oxidative metabolism in mitochondria, which is dependent on lipid utilization ^27,28^. This transition is part of the establishment of more efficient energy production systems, to face the substantial increase in cardiac workload resulting from the growth of the organism ^28,29^.

Our data suggest that GR controls the expression of genes involved in postnatal metabolic maturation, including those involved in fatty acid oxidation, mitochondrial respiration and energy transfer from mitochondria to the cytosol.

Gene ontology analysis according to cellular components (**Fig. 5e**), suggested again that significant changes occur in gene networks connected with mitochondrial electron transfer flavoprotein complex (**Supplementary Figure 6a**) and highlighted major differences in genes codifying for contractile fiber proteins (**Supplementary Figure 6b**), in particular in proteins participating to the I band and Z disk (**Fig. 5f**). These data suggest that GR regulates the cytoarchitectural structure of cardiomyocytes.

During the early postnatal development, the cytoarchitectural structure of cardiomyocytes shifts from loose spatial organization to highly organized and more efficient sarcomere units ^28,30^. To evaluate a potential impact of GR ablation on this physiological reorganization, we performed ultrastructure analysis by transmission electron microscopy of GR-cKO and control cardiomyocytes at birth and during the early postnatal development. As well documented in the literature for neonatal cardiomyocytes, P1 control cardiomyocytes displayed an immature myofibrillar architecture characterized by few and chaotically-arranged myofibrils, and large free cytoplasmic space between myofibrils and isolated mitochondria (**Fig. 6a**, mitochondria magnification in **6c)**. Neonatal GR-cKO cardiomyocytes appeared slightly more immature, as visible by less numerous myofibrils, randomly organized with cellular substructures, sometimes arranged around irregularly-shaped nuclei (**Fig. 6b**, mitochondria magnification in **6d)**. These data are in line with a previously reported role for GR in late gestation foetal cardiomyocyte maturation ^18,19,22,23^.

**Figure 6.**
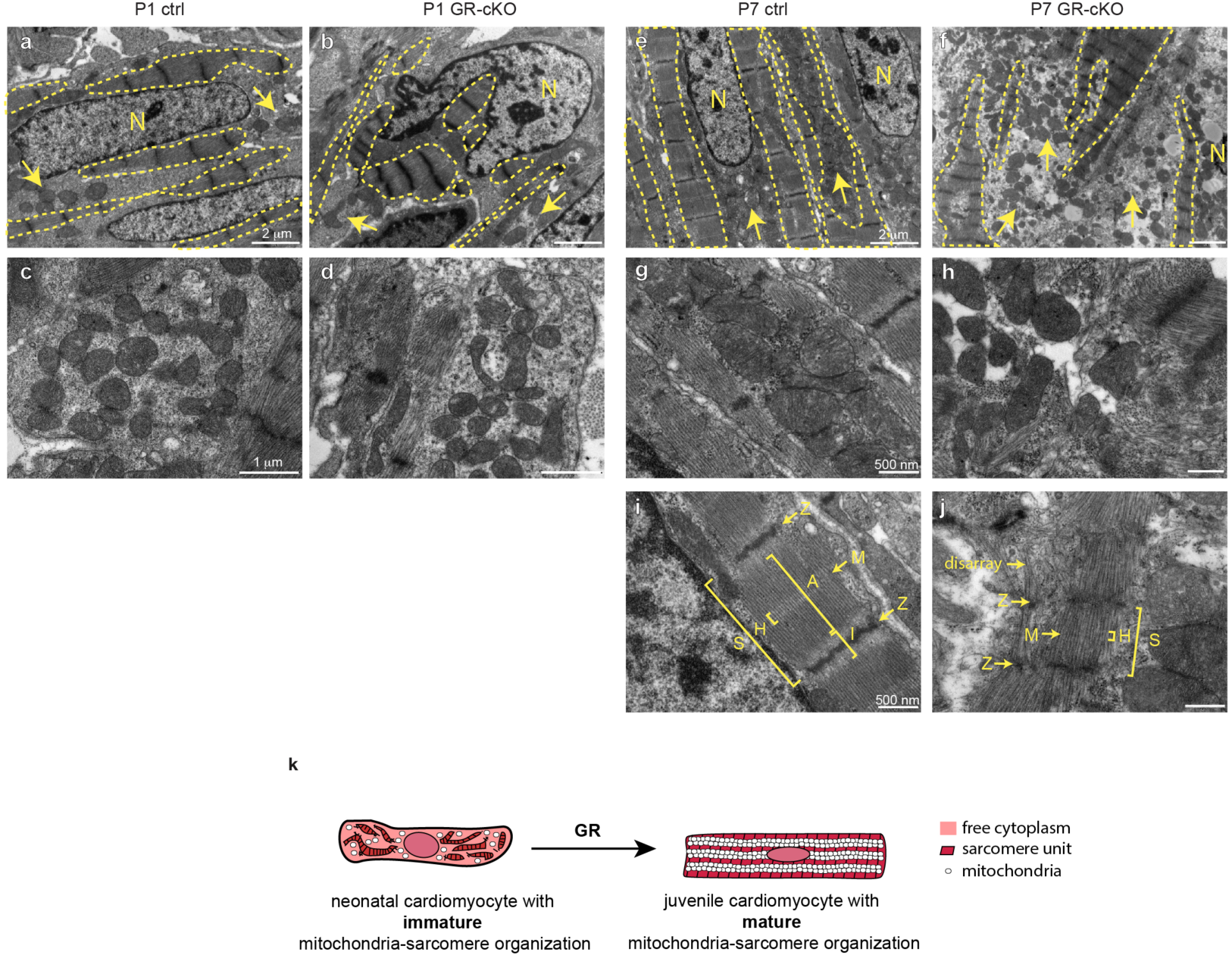
GR-cKO inhibits early post-natal cardiomyocyte maturation *in vivo*. (**a-d**) Electron microscopy analysis of P1 control (**a, c**) and GR-cKO (**b, d**) heart sections, showing a slight more immature myofibrillar architecture in GR-cKO cardiomyocytes, characterized by less numerous myofibrils, randomly organized with cellular substructures (dotted lines **in a-b**) and isolated mitochondria in both (arrows in **a, b** and magnification in **c, d**); Electron microscopy analysis of P7 control (**e, g, i**) and GR-cKO (**f, h, j**) heart sections: control cardiomyocytes show organized myofibrillar architecture (dotted lines in **e**, magnification in **i**) in close contact with clusters of mitochondria (arrows in **e**, magnification in **g**), whereas GR-cKO cardiomyocytes show irregular and immature myofibrils (dotted lines in **f**, magnification in **j**) arranged around isolated not organized mitochondria (arrows in **f**, magnification in **h**); (**k**) Schematic diagram showing the role of GR in promoting maturation of sarcomere-mitochondrial organization in cardiomyocytes during the early postnatal development; N = nucleus; S = sarcomere unit; H = H band; A = A band; I = I band; M = M band; Z = Z line.

One week later, at post-natal-day-7 (P7), control cardiomyocytes had substantially little free cytoplasmic space, with aligned and organized myofibrils, which appear in close contact with clustered mitochondria (**Fig. 6e**, mitochondria magnification in **6g**). Myofibrils appeared regularly and parallel oriented with normal Z line banding (**Fig. 6i**). These observations are in line with previously documented cardiomyocyte cytoarchitecture during early post-natal maturation, which is coupled with the building-up of energy micro-domains, responsible for a more efficient energy transfer system from mitochondria to sarcomere structures ^31^. By contrast, at this stage, GR-cKO cardiomyocytes retained multiple features that characterize immature neonatal/foetal cardiomyocytes, including a high content of free cytoplasm with an irregular pattern of myofibrils, often misaligned, thus losing the major axis orientation, and arranged around isolated mitochondria (**Fig. 6f**, mitochondria magnification in **6h**). Furthermore, the sarcomere apparatus of GR-cKO at this stage was less mature than controls, as visible by more disorganized sarcomere architecture, featuring a less evident A, I and H bands and M line (**Fig. 6j**). Immunofluorescence analysis by confocal microscopy of myofibrillar structures on heart sections from P7 GR-cKO compared to control mouse model confirmed the presence of cardiomyocytes with reduced and partially disassembled myofibrils (**Supplementary Fig. 7a**).

Altogether, our data show that mitochondria-myofibrils organization at the age of postnatal day 7 in GR-cKO mice resembles the organization of neonatal cells, indicating that GR plays a key role in the maturation of mitochondria-myofibrils organization occurring in the early postnatal period (**Fig. 6k**).

### Antagonising glucocorticoid receptor promotes cardiomyocyte proliferation and heart regeneration after myocardial infarction

All the data collected so far strongly supported a role of GCs/GR axis activation in promoting maturation, thus restraining the proliferation of cardiomyocytes in early postnatal development. Next, we tested whether GR ablation is sufficient to promote cardiomyocyte proliferation upon myocardial injury. To this end, we induced myocardial infarction (MI) in P7 GR-cKO and control mice by permanent ligation of the left anterior descending coronary artery, and we evaluated cardiomyocyte proliferation 10 days post-MI (**Fig. 7a**). Cardiomyocyte cell cycle re-entry was increased both in remote and border zones of myocardial infarction in GR-cKO mice (**Fig. 7b**). Furthermore, the level of cycling cardiomyocytes with marginalization of sarcomeric structures to the cell periphery was robustly increased in GR-cKO hearts compared to controls (**Fig. 7c)**. This phenomenon has been documented in the entire heart during the spontaneous regeneration process following cardiac injury in neonatal mice and it is considered as a prerequisite to effective cardiomyocyte division ^5,7,30,32–34^. Furthermore, Aurora B positive cardiomyocytes were increased in the border zone in GR-cKO hearts (**Fig. 7d**). These data, therefore, suggest that GR-cKO cardiomyocytes after myocardial infarction are more prone to disassemble sarcomeric components in order to undergo cell division.

**Figure 7.**
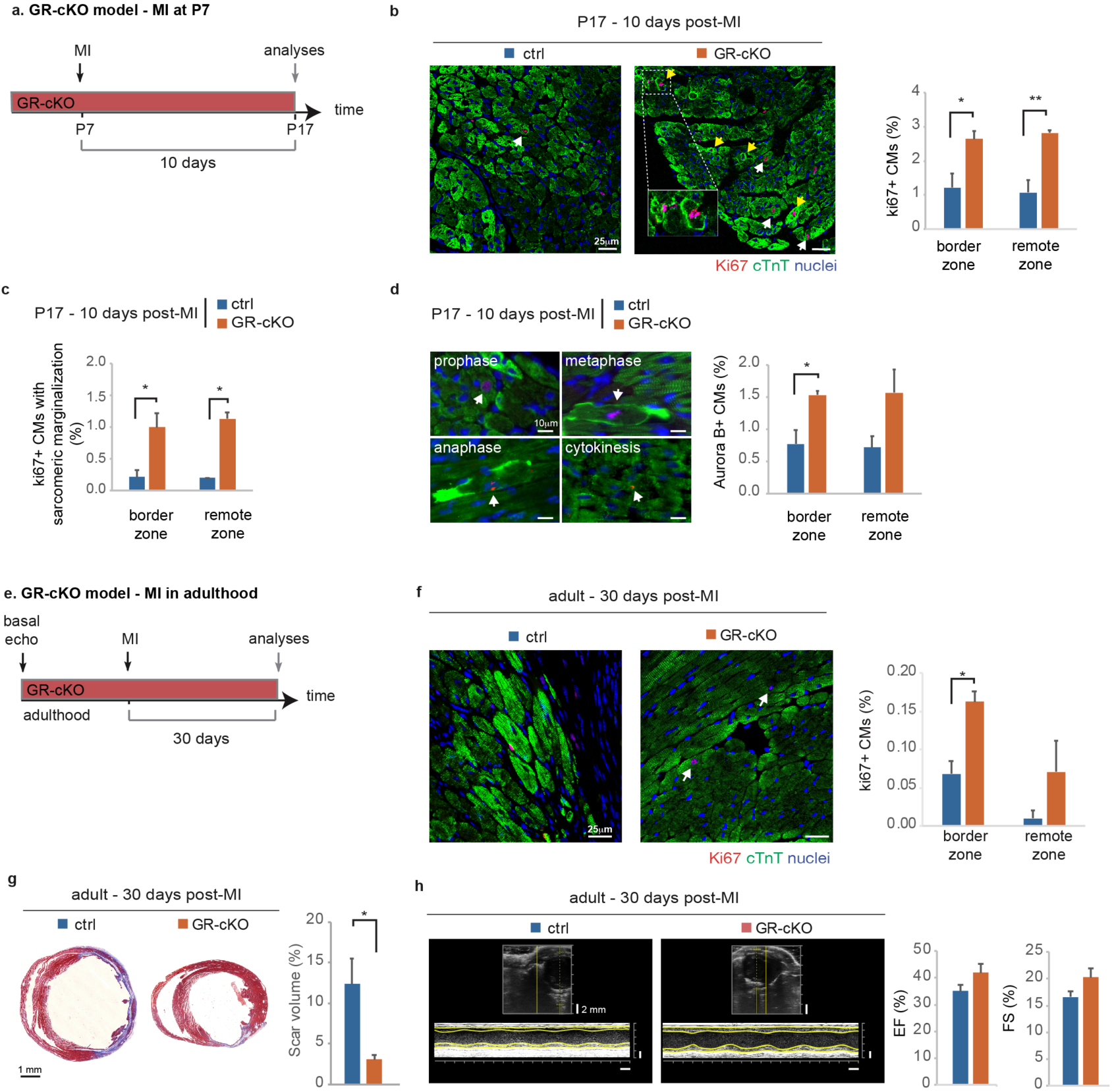
GR ablation enhances mammalian cardiac regenerative ability. **(a)** Experimental design for analysis of cardiomyocyte dedifferentiation and proliferation following injury at juvenile (Postnatal-day-7 - P7) stage in control and GR-cKO mice; (**b-c**) *in vivo* evaluation of juvenile cardiomyocyte (CM) cell-cycle re-entry (ki67+, **b**) and sarcomeric marginalization (**c**) by immunofluorescence analysis of Ki67 and cardiac Troponin T in control and GR-cKO heart sections 10 days following MI according to the scheme in **a**; representative pictures of cell-cycle activity and marginalization of sarcomeric structures in the border zone of ctrl and GR-cKO cardiomyocytes are provided; yellow and white arrows point at proliferating cardiomyocytes with and without sarcomere marginalization, respectively; scale bars, 25 µm; (**d**) *in vivo* evaluation of juvenile Aurora B kinase positive CMs; representative pictures are provided; scale bars, 10 μm; **(e)** Experimental design for cardiac regeneration analysis following injury in adult (2-months-old) control and GR-cKO mice; (**f**) *in vivo* evaluation of adult cardiomyocyte cell-cycle re-entry (ki67) by immunofluorescence analysis of Ki67 in control and GR-cKO heart sections ∼1 month following MI according to the scheme in **e**; **(g**) Scar quantification based on Masson’s trichrome staining of heart sections of control and GR-cKO mice ∼1 month post-MI according to the scheme in **e**; representative pictures are provided; scale bars, 1 mm; (**h**) Echocardiographic measurements of ejection fraction (EF) and fractional shortening (FS) of injured control and GR-cKO adult mice ∼1 month post-MI according to the scheme in **e**; representative images of M-mode echocardiographic analyses are provided; scale bars, 2 mm (vertical axis) and 200ms (horizontal axis); In all panels numerical data are presented as mean (error bars show s.e.m.); * p < 0.05, ** p < 0.01, *** p < 0.001 and **** p < 0.0001.

In order to evaluate GR suppression as a potential regenerative strategy in adult mammals, we analysed cardiomyocyte proliferation and cardiac tissue replacement following myocardial infarction in GR-cKO and control adult mice (**Fig. 7e**). Consistent with the juvenile settings, cardiomyocyte proliferation was increased in GR-cKO compared to control adult hearts in the infarct border zone (**Fig. 7f**). In addition, GR-cKO adult hearts displayed reduced scarring (**Fig. 7g**) and a mild overall improvement in cardiac performance (**Fig. 7h**). These data demonstrate that the deletion of GR facilitates cell cycle re-entry and proliferation upon cardiac injury, resulting in tissue replacement with reduced scar formation.

Collectively, our findings indicate that GR ablation facilitates heart regeneration in adult mice after myocardial infarction as a result of increased cardiomyocyte dedifferentiation and proliferation.

## DISCUSSION

Our findings unveiled an important role for GCs/GR axis in promoting cardiomyocyte maturation while restraining the proliferative potential during physiological postnatal cardiac development, and following injuries in juvenile and adult stages (**Fig. 8**). First, our data show that endogenous glucocorticoids, namely corticosterone in the mouse model, potently suppress the proliferation of immature neonatal cardiomyocytes. These findings highlight the need for a careful evaluation of cardiomyocyte renewal in conditions characterized by high levels of endogenous glucocorticoids, for example, upon prolonged stress or in Cushing syndrome. Mice with global inactivation of the GR gene die soon after birth, principally due to impaired lung maturation, though effects on the liver are also critical ^22^. GR is detectable in foetal heart starting from mid-gestation, but it exerts its activity at late gestation when circulating glucocorticoid levels increase ^22^. Our data are consistent with previous studies suggesting that GR promotes the maturation of foetal cardiomyocytes ^22,35,36^. Notably, we expand this knowledge by demonstrating that a physiological increase in GCs/GR axis activation in the early postnatal development is responsible for cardiomyocyte maturation and loss of proliferative ability. In this regard, we observed a cardiomyocyte-specific increase in GR abundance during the early postnatal development, consistent with the loss of postnatal cardiac regenerative ability. An *in vivo* mouse model for cardiomyocyte-specific GR ablation (GR-cKO) has been previously generated and its heart morphology and function has been reported to appear normal till adulthood, although a decline in heart function starting at 3 months of age and premature death for heart failure later on has been documented ^26^. By careful evaluation of this *in vivo* model during the early post-natal development, we here unveiled that cardiomyocytes lacking GR undergo reduced cell cycle exit and hypertrophic growth. We also observed that cardiomyocytes lacking GR display a less mature transcriptional signature and ultrastructural morphology, with reduced expression of genes involved in the transition towards fatty acid oxidation, mitochondrial respiration and energy transfer from mitochondria to the cytosol, along with an impairment in the building-up of a mature myofibrils-mitochondria organization. We thus propose that postnatal GCs/GR axis activation takes part in the hyperplastic to hypertrophic transition as well as in the metabolic and cytoarchitecture maturation occurring shortly after birth.

**Figure 8.**
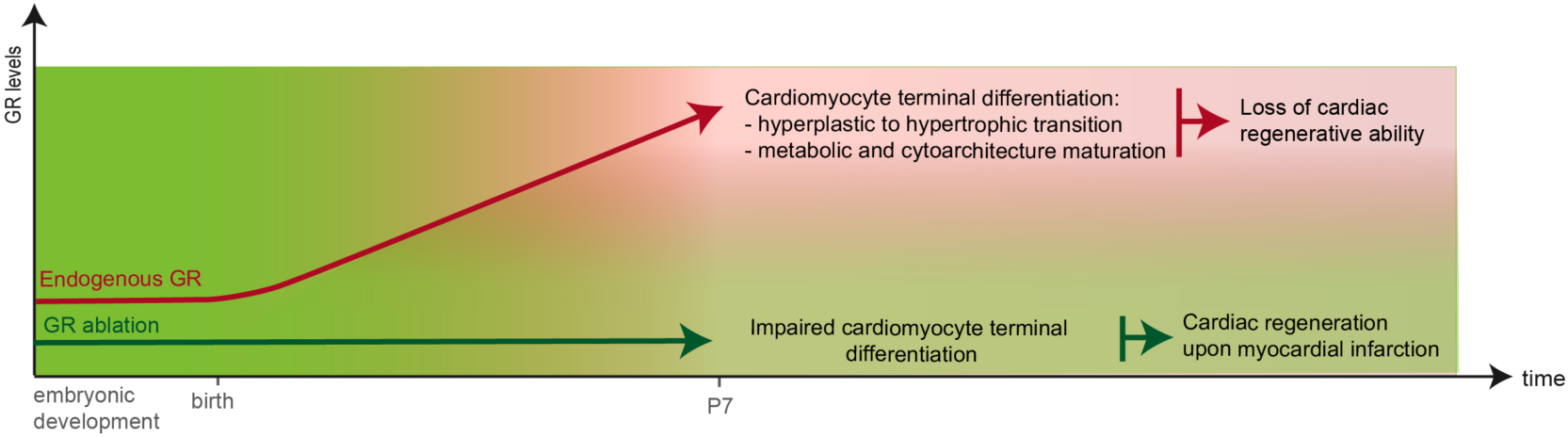
GR inhibition facilitates cardiomyocyte proliferation and broadens the postnatal regenerative window in the mouse model. A schematic diagram of the role of GR in controlling postnatal cardiomyocyte proliferation and heart regenerative ability. A physiological increase of GR levels (red line) during the first week of postnatal life promotes cardiomyocytes cell-cycle exit, hypertrophic growth and mitocondria-sarcomere maturation, thus resulting in loss of regenerative ability. GR ablation (green line) in cardiomyocytes impairs cardiomyocytes terminal differentiation and facilitate cardiomyocytes proliferation and cardiac regeneration following injury.

Intriguingly, reduced mitochondrial gene expression and decreased oxidative phosphorylation activity have been recently observed in proliferating cardiomyocytes in regenerating heart models ^37^. Additionally, sarcomere disassembly is a pre-requisite to cardiomyocyte proliferation in models of spontaneous cardiac regeneration, such as zebrafish and neonatal mice ^5,7,30,32–34^, and reduced sarcomere expression and organization in cardiomyocytes has been linked to increased ability to proliferate ^30,38,39^. Thus, both metabolic and sarcomere immaturity of GR-cKO cardiomyocytes likely facilitate their proliferation. Intriguingly, our data point at proteins of the I band and/or Z-disk, where the first step of sarcomere disassembly in proliferating cardiomyocytes has been previously reported ^32^. Among downregulated I band genes in GR-cKO we observed PDLIM5 (PDZ and LIM domain protein 5, also known as Enigma Homolog - ENH), which plays a key role in Z-line structure ^40^; PGM1 (Phosphoglucomutase 1), an enzyme that is involved in energy metabolism, whose reduction lead to increased flow of glucose into glycolysis ^41^, and may localize at the Z disk in stress conditions ^42^; ANK2 (Ankyrin-B), which is involved in calcium handling ^43^; RYR3 (Ryanodine Receptor type 3), which is mainly involved in calcium cycling in purkinje fibers ^44,45^; MYZAP (Myocardial Zonula Adherens Protein), which is a component of the intercalated disk important for sustaining heart function ^46^. Thus, GR regulates the expression of genes involved in the cytoarchitectural structure of cardiomyocytes connected to both energy metabolism and calcium handling. The immaturity of GR-cKO cardiomyocytes may also explain the reported decline in heart function starting at 3 months of age and the premature death for heart failure observed in these mice ^26^. In this regards adult GR-cKO hearts were reported to have reduced levels of Ryanodine receptor (RyR2) and impairment in calcium handling, which resulted in contractile dysfunction ^26^. Our data suggest that this phenotype could be placed into a wider framework of incomplete cardiomyocyte maturation.

Notably, GR ablated cardiomyocytes were facilitated to re-enter into the cell cycle, disassembly their sarcomere structures and divide upon cardiac injuries in both juvenile and adult mice, thus promoting cardiac regeneration with reduced scar formation and improved muscle tissue replacement. Therefore, we suggest that GR antagonization could be employed as a novel approach for heart regeneration based on endogenous cardiomyocyte renewal. Previous studies demonstrated that GR restrains the regeneration of skeletal muscles ^47^ and the zebrafish fin ^48^. Considering the systemic action of circulating glucocorticoids, the unveiled role of GR as a “brake” for cardiac regenerative ability could be placed into a wider framework of GR as a restrainer of the regenerative ability of multiple organs.

Clinically, glucocorticoids are employed in the treatment of many inflammatory conditions, exerting their primary action in suppressing inflammatory immune cells activity ^49^. We here unveiled that synthetic glucocorticoids routinely used in clinical settings are able to reduce cardiomyocyte proliferative potential. This is in line with recent findings suggesting that dexamethasone promotes terminal differentiation of neonatal cardiomyocytes *in vitro* ^50^ and reduces cardiomyocyte proliferation during post-natal development *in vivo*, resulting in a decreased cardiomyocyte endowment in the developing heart ^51,52^. In addition, a significant association between the administration of glucocorticoids and heart failure has been reported in epidemiological studies ^53,54^. Our study may explain this association as a consequence of the reduction of cardiomyocyte turnover, a process that was recently reported in adult mammalian hearts, including humans ^10,11^. These findings highlight the importance of careful scheduling of these agents in clinical settings, especially during the postnatal developmental period.

The possibility to interfere with inflammatory processes post-MI by the administration of glucocorticoids has been studied for a long time in the attempt to reduce fibrotic remodelling ^55^. However, inflammation is also a required factor to achieve regeneration of multiple organs ^56^, including the heart of neonatal mice following injury ^57,58^. The results of studies analysing the effects of glucocorticoid delivery after myocardial infarction are controversial, with some studies reporting an increase in the incidence of left ventricular free wall rupture and patients death ^59–61^, while others showing a reduction of infarct size and improved heart function ^62–64^. Our findings suggest that the activation of GR may support heart contractility by preserving cardiomyocyte maturity, however inhibiting sarcomere disassembly and suppressing the generation of new cardiomyocytes. Thus, the overall balance between heart function and regeneration as a consequence of insufficient or excessive GR activity could likely explain the reported discrepancies.

Most studies on GR were focused on immune cells or other stromal cardiac components, thus surprisingly little is known on the effect of GR activity in cardiomyocytes. Our transgenic strategy, using a cardiomyocyte-specific promoter (Myh6) to ablate GR expression, unveiled a direct effect of glucocorticoid signalling in cardiomyocytes, influencing their proliferative potential and regenerative response.

In summary, our study provides important insights into the postnatal activity of GCs/GR axis in cardiomyocyte maturation and cell cycle exit, and suggests a rationale for boosting the proliferative ability of adult cardiomyocyte for inducing cardiac regeneration.

## ACKNOWLEDGMENTS

This project has received funding from the European Union’s Horizon 2020 research and innovation programme under the ERA-NET Co-fund action N°680969 to Gabriele D’Uva and Eldad Tzahor, from Fondazione Cariplo (Grant Number: GR 2017-0800 to Gabriele D’Uva and Mattia Lauriola), and from International Society for Heart Research (ISHR research fellowship 2019 to Nicola Pianca).

We would like to thank Prof. Giuseppe Pelosi, Dr. Donata Micello, Emanuela Paiola, Cristina Kluc, Alessandra Perlasca and Barnaba Rainoldi for technical assistance in sectioning of paraffin-embedded samples. Part of this work was carried out in ALEMBIC, an advanced microscopy laboratory established by IRCCS Ospedale San Raffaele and Università Vita-Salute San Raffaele. We would like to thank Desirè Zambroni for the technical assistance in the *in vitro* immunofluorescence imaging.

## METHODS AND METHODS

### Mouse experiments

Experiments were approved by the Animal Care and Use Committee of the University of Bologna and Weizmann Institute of Science. Cardiomyocyte-specific GR knock-out (GR-cKO) mice were generated by mating animals expressing Cre under the promoter of Myh6 with animals harbouring loxP sites flanking the exon 3 and 4 of GR gene, as previously reported ^26^ (see **Fig. 3a**). Oligonucleotide sequences for genotyping the above-described mouse lines are listed in **Supplementary Table 1**.

### Myocardial infarction

Myocardial infarction at juvenile (P7) or adult stage was induced by ligation of the left anterior descending coronary artery as previously described ^38,65^. P7 mice were anaesthetized by cooling on an ice bed for 4 min, whereas adult mice were sedated with isoflurane (Abbott Laboratories) and, following tracheal intubation, were artificially ventilated. Lateral thoracotomy at the fourth intercostal space was performed by blunt dissection of the intercostal muscles following skin incision. Following ligation of the left anterior descending coronary artery, thoracic wall incisions were sutured with 6.0 non-absorbable silk sutures, and the skin wound closed using skin adhesive. Mice were then warmed for several minutes until recovery. The hearts were collected for analysis at different time points, as indicated in **Fig. 7**.

### Echocardiography

Heart function was evaluated by transthoracic echocardiography performed on sedated mice (isoflurane, Abbott Laboratories) using a VisualSonics Vevo 3100^®^ or MyLab 70 XV devices.

### Histology (Masson’s trichrome)

Murine hearts were fixed in 4% paraformaldehyde for 24 hours, then embedded in paraffin, sectioned (3μm thick) with a microtome (Leica) and attached to SuperFrost II plus slides (Thermo). For analysis of cardiac regeneration following the myocardial infarction procedure, paraffin sections were cut through the entire ventricle from apex to base into serial sections. Masson’s trichrome staining was used for the detection of fibrosis and performed with a standard kit (Bio-Optica, 04-010802A) according to the manufacturer’s protocol. Scar size was quantified in each section with ImageJ software (National Institutes of Health) based on Masson’s trichrome staining. The percentage of myocardial tissue volume occupied by scar was calculated as the sum of scar volumes between serial sections normalized on the total left ventricular and septal tissue volume. The volume between serial sections was calculated with the formula of a truncated pyramid: 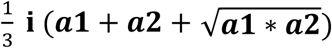, where **a1** and **a2** are the areas of 2 serial sections and **i** is the interval (0.2-0.4 mm) between them.

### Cardiomyocyte isolation and culture

Primary neonatal cardiomyocytes were extracted from hearts of 0-day-old or 1-day-old (P0-P1) mice. Cardiac cells were isolated by enzymatic digestion with pancreatin and collagenase, as previously described ^66^. Cells were then cultured in gelatine-coated (0.1%, G1393, Sigma) wells with DMEM/F12 (Aurogene) supplemented with L-glutamine, sodium pyruvate, non-essential amino acids, penicillin, streptomycin, 5% horse serum and 10% FBS (hereafter referred to as ‘complete-medium’) at 37°C and 5% CO_2_. The cells were allowed to adhere for 48 h in complete-medium. Subsequently, the medium was replaced with an FBS-deprived complete-medium containing Corticosterone at the indicated concentrations (27840, Sigma) for 48 hours. For BrdU assay, BrdU (10 µM, B5002, Sigma) was introduced along with the treatments.

### Cardiomyocyte and stromal cell separation

To assess the differential gene expression in cardiomyocytes and stromal cells in P1 and P7 hearts, we proceeded as follows. P1 hearts were subjected to enzymatic digestion as described above, whereas P7 hearts were harvested and anterogradely perfused with digestion enzymes (collagenase, trypsin, protease) as previously described in ^67^. Cardiomyocytes were separated from stromal cells using the immuno-magnetic cell sorting MACS Neonatal Isolation System (130-100-825, Miltenyi biotech), following the manufacturer’s protocol. The different cells were then centrifuged, and total RNA was extracted from the pellet as described below.

### Immunofluorescence on cells and tissue sections

Cultured cells were fixed with PFA 4% (Sigma, diluted in PBS) for 20 minutes at 4°C. Heart sections underwent deparaffinization (by immersion in Xylene and rehydration by immersion in solutions with a decreasing concentration of Ethanol), and heat-induced antigen retrieval in a sodium citrate solution (Sigma) at pH 6.0, followed by gradual chilling. Then heart sections and cultured cells were processed in the following manner. Samples were permeabilized with 0.5% Triton-X100 (Sigma) in PBS for 5 minutes at room temperature and the aspecific binding of the antibodies was prevented by applying a blocking solution (PBS supplemented with 5% BSA (Sigma) and 0.1% Triton-X100) for 1 hour at room temperature. For the BrdU staining protocol, a DNA hydrolysis step between the permeabilization and the blocking step was performed by the addition of 2M HCL (Sigma) for 30 minutes at +37°C and then 3 washes in PBS.

Then samples were incubated overnight at 4° C with the following antibodies diluted in PBS supplemented with 3% BSA and 0.1% Triton-X100. Anti-cTnT (1:200, ab33589, Abcam) and anti-cTnI (1:200, ab47003, Abcam) antibodies were used to identify cardiomyocytes. Anti-Ki67 antibody (1:100, ab16667, Abcam), anti-BRDU antibody (1:50, G3G4, DSHB), and anti-aurora B (1:50, 611082, BD Transduction Laboratories) antibodies were used to analyse cell-cycle re-entry, DNA synthesis and cytokinesis, respectively. Other antibodies and dyes used in the study: anti-GR (1:50,12041S, Cell Signalling), wheat germ agglutinin (WGA) AlexaFluor488-conjugated (1:200, Invitrogen, 29022-1).

After primary antibody incubation, 3 washes in PBS were performed and samples were incubated for 1 hour at room temperature with fluorescent secondary antibodies, diluted in PBS supplemented with 1% BSA and 0.1% Triton-X100. Secondary antibodies used: anti-mouse AlexaFluor 488 (115-545-003, Jackson), anti-rabbit AlexaFluor 488 (111-545-003, Jackson), anti-mouse Cy3 (115-165-003, Jackson), anti-rabbit 594 (AlexaFluor 111-585-003, Jackson).

After 3 washes in PBS, DAPI (4’,6-diamidino-2-phenylindole dihydrochloride, Sigma), diluted in PBS, was applied for 15 minutes at room temperature for nuclei visualization. Samples were then washed 2 more times in PBS. Cells in culture plates were imaged at a widefield microscopy Zeiss (Axio Observer A1) or at a widefield microscopy ArrayScan XTI (ThermoFisher). Slides were mounted with an antifade solution (Vectorlabs), covered with a coverslip, sealed with nail polish and imaged at a widefield fluorescent microscopy Zeiss (Axio Observer A1), at a widefield fluorescent microscope (Nikon Eclipse 80i) and at a confocal fluorescent microscope (Leica TCS SP5).

### Assessment of cardiomyocyte size

The cross-sectional area of cardiomyocytes in P7 hearts was assessed using ImageJ software (National Institutes of Health) on the basis of wheat germ agglutinin (WGA) staining. For that, slides were deparaffinized as described above, rinsed in PBS and then incubated overnight at +4°C with WGA conjugated to Alexa Fluor 488 (1:200, Invitrogen, 29022-1). Slides were then rinsed 3 times in PBS, mounted with an antifade solution (Vectorlabs) and imaged at widefield microscope (Nikon Eclipse 80i). Only cardiomyocytes that were aligned transversely were considered for the quantification of the cross-sectional area.

### Protein extraction and Western Blotting

Western blotting was performed with the SDS–PAGE Electrophoresis System. Proteins were extracted with RIPA buffer with the addition of proteinase inhibitor (Sigma) and phosphatase inhibitors (cocktail 2 and 3, Sigma). SDS-PAGE was performed with 60μg of protein extracts and transferred to PVDF membrane, which was then incubated with the following antibodies: anti-GR (12041S, Cell Signalling) and anti-GAPDH (2118, Cell Signalling). Horseradish peroxidase (HRP) conjugated anti-rabbit antibody (NA934, GE) was used as secondary antibody. The signal was detected by LICOR imaging system with ECL substrate (LICOR).

### Transcriptional analysis

Total RNA extraction was performed with NucleoSpin RNA II kit (Macherey Nagel) according to the manufacturer’s protocol. RNA quantification and quality check were performed using a Nanodrop spectrophotometer (N1000, Thermo). RNA was reverse transcribed to cDNA with VILO (Thermo Fisher) according to the manufacture’s protocol. Real Time (rt)-PCR was performed using Fast SYBR Green PCR Master Mix (Applied Biosystems) on a QuantStudio 6 Flex instrument (Applied Biosystems). Oligonucleotide sequences for real-time PCR analysis performed in this study are listed in **Supplementary Table 2**.

### RNA sequencing

Transcriptomic analysis was performed using QuantSeq FWD 3’ mRNA-Seq Services of Lexogen. Briefly, purified total RNA was extracted from in vitro cultures of cardiomyocytes and from whole heart homogenates; sequencing was performed on NextSeq 500 (Illumina) and quantification was based on tags in 3’ region with more than 20M reads per sample; data analysis for differential expression was conducted with DESeq2 (Bioconductor) and with Galaxy (https://usegalaxy.org/).

### Electron Microscopy

Small myocardial samples were fixed after surgery in 2.5% glutaraldehyde in 0.1 M cacodylate buffer at pH 7.4 and postfixed in 1% OsO_4_ in the same buffer. After dehydration in graded ethanol specimens were embedded in Araldite. Thin sections were stained in uranyl acetate and lead citrate and observed in a CM10 transmission electron microscope.

### Statistical analysis

Statistical analyses were performed with GraphPad software (Prism). Whenever normality could be assumed the Student t-test or analysis of variance (ANOVA) followed by Tukey’s or Sidak’s test was used to compare group means, as specified in the figure legends. P value<0.05 was considered to represent a statistically significant difference. In all panels, numerical data are presented as mean+s.e.m.; results are marked with one asterisk (*) if P <0.05, two (**) if P <0.01, three (***) if P <0.001 and four (****) if P <0.0001;

