## Supplementary data for "Glucocorticoid Receptor ablation promotes cardiac regeneration by hampering cardiomyocyte terminal differentiation"

### SUPPLEMENTARY FIGURES AND TABLES

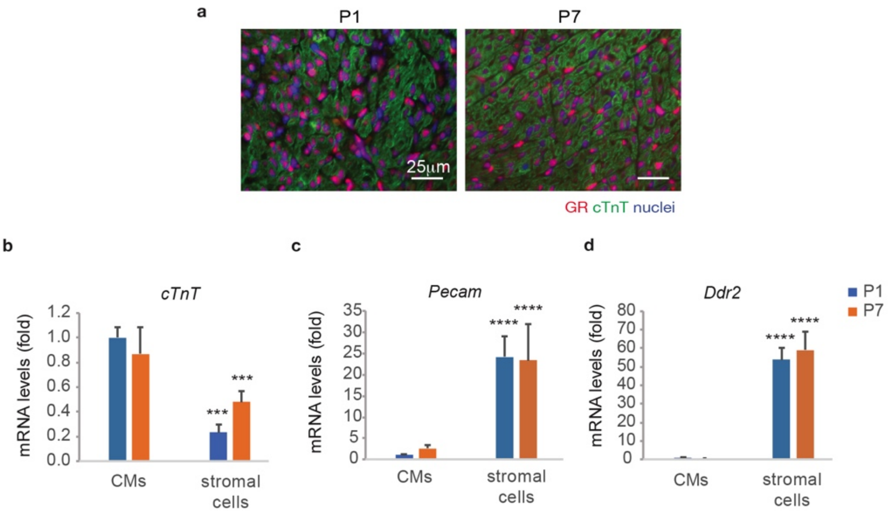


**Supplementary Figure 1. GR nuclear localization and validation of separation between cardiomyocyte and stromal cells preparations. (a**) Immunofluorescence analysis of GR in P1 and P7 left ventricular heart sections, showing similar nuclear localization of GR in different cell types; scalebar = 25 µm; (**b-d**) Cardiomyocytes (CMs) and stromal cells isolated from P1 and P7 hearts were analysed by RT-PCR for markers of the major cardiac cell types, namely cardiomyocytes (**b**, cardiac troponin T), endothelial cells (**c**, Pecam) and fibroblasts (**d**, Ddr2), showing the absence of other cell types markers in cardiomyocytes preparations. In all panels, numerical data are presented as mean (error bars show s.e.m.); *** $p < 0.001$ and **** $p < 0.0001$.

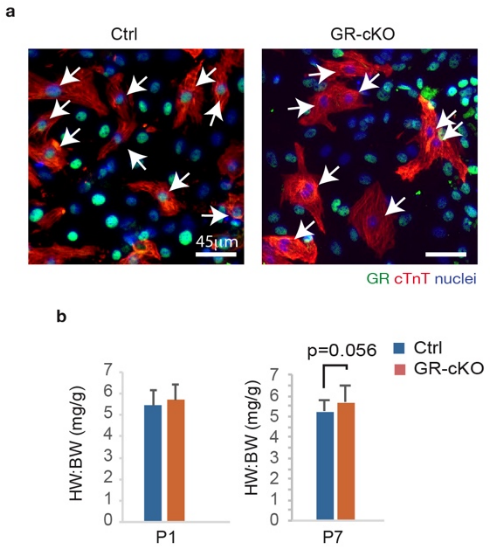


**Supplementary Figure 2.** ***In vitro*** **and** ***in vivo*** **analysis of cardiomyocyte-specific GR Knock Out (GR-cKO) mouse model. (a)** Immunofluorescence analysis of GR in cardiomyocyte cell cultures isolated from postnatal-day-1 (P1) GR-cKO and control mice, showing the absence of GR expression in cardiomyocytes isolated from GR-cKO mice; arrows point at GR-positive and GR-negative nuclei in controls and GR-cKO cardiomyocytes, respectively; **(b)** Heart weight (HW) to body weight (BW) ratio in P1 and P7 ctrl and GR-cKO mice.

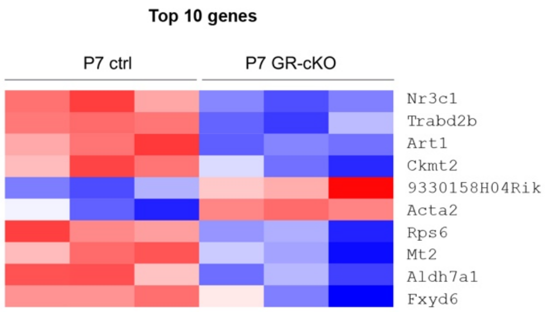


**Supplementary Figure 3. Postnatal-day-7 Gr-cKO vs ctrl hearts: TOP Ten genes by p-value.** Heatmap of top ten differentially expressed genes by RNA-Seq transcriptome analysis of P7 GR-cKO versus control hearts.

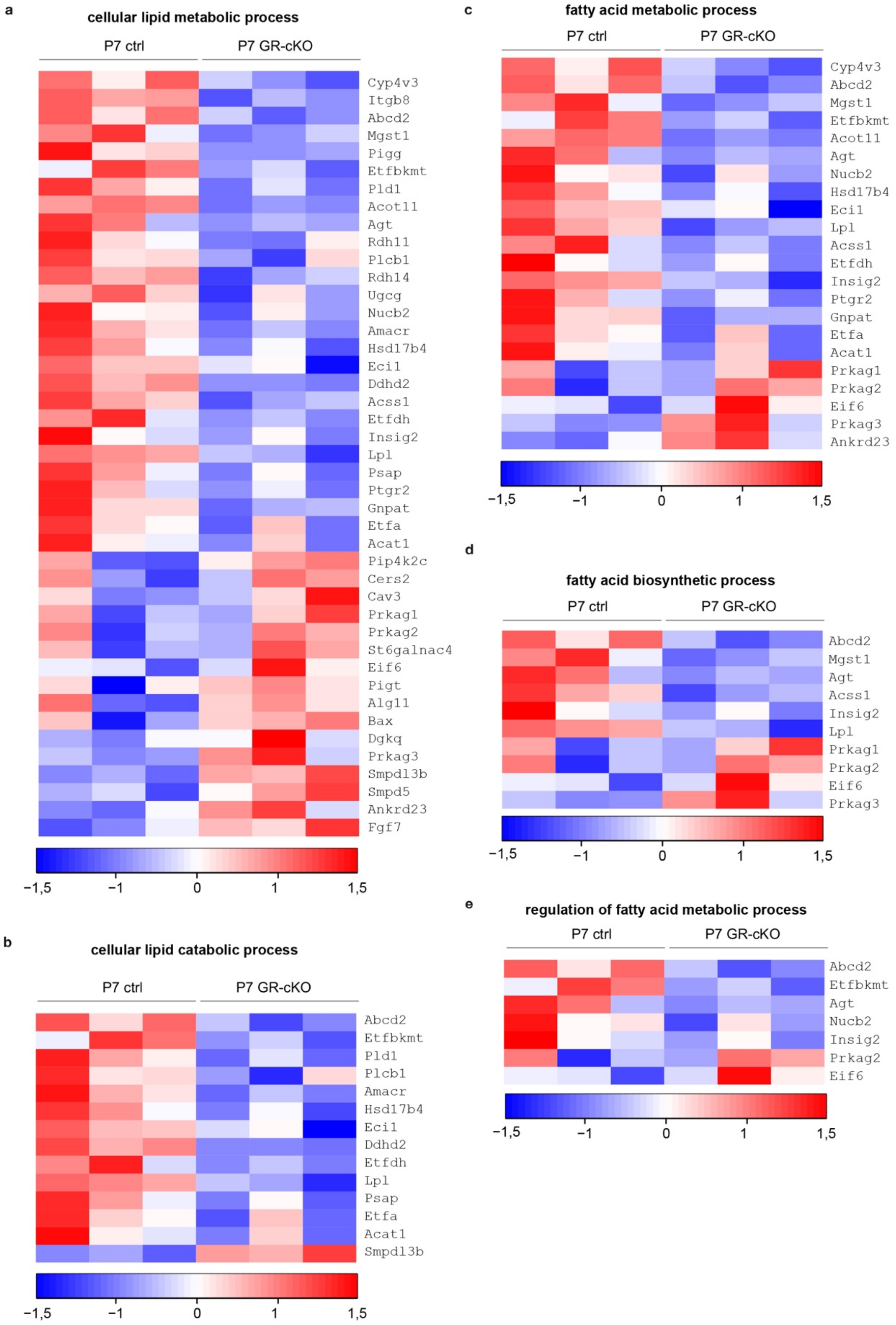


**Supplementary Figure 4. Heat maps of gene ontology (GO) analysis of postnatal-day-7 GR-cKO versus controls hearts according to Biological Process. (a-e)** Heat maps of differentially expressed genes by RNA-Seq transcriptome analysis of P7 GR-cKO versus controls hearts according to the following Biological Process gene ontology terms: lipid metabolic process (GO:0044255, **a**), fatty acid metabolic process (GO:0006631, **b**), fatty acid biosynthetic process (GO:0006633, **c**), regulation of fatty acid metabolic process (GO:0019217, **d**) and cellular lipid catabolic process (GO:0044242, **e**).

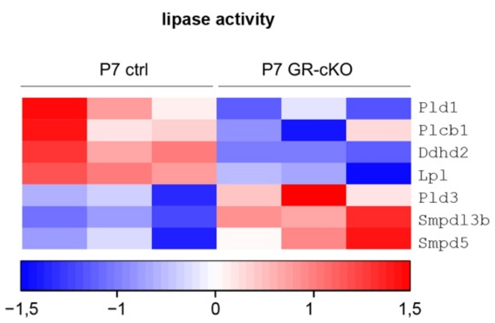


**Supplementary Figure 5. Heat maps of gene ontology (GO) analysis of postnatal-day-7 GR-cKO versus controls hearts according to Molecular Function.** Heat map of differentially expressed genes by RNA-Seq transcriptome analysis of P7 GR-cKO versus controls hearts according to lipase activity (GO:0016298).

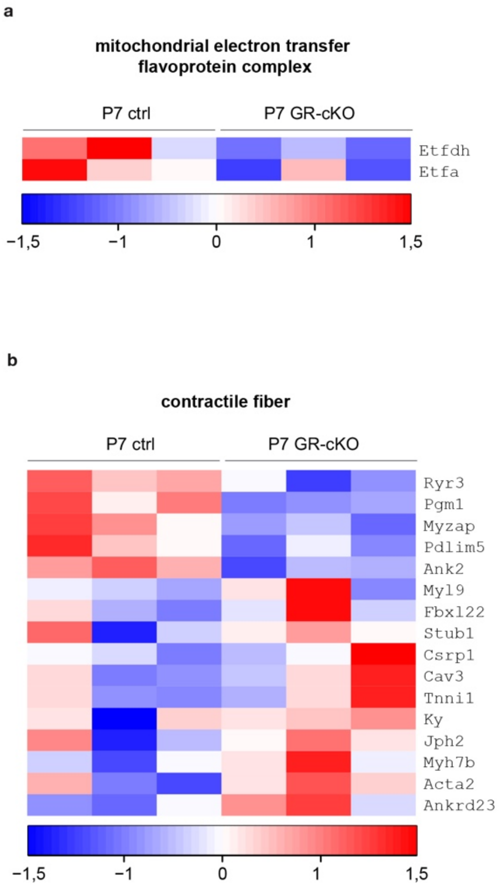


**Supplementary Figure 6. Heat maps of gene ontology (GO) analysis of postnatal-day-7 GR-cKO versus controls hearts according to Cellular Components.** (**a,b**) Heat maps of differentially expressed genes by RNA-Seq transcriptome analysis of P7 GR-cKO versus control hearts according to the following Cellular Components gene ontology terms: mitochondrial electron transfer flavoprotein complex (GO:0017133, **a**) and contractile fiber (GO:0043292, **b**).

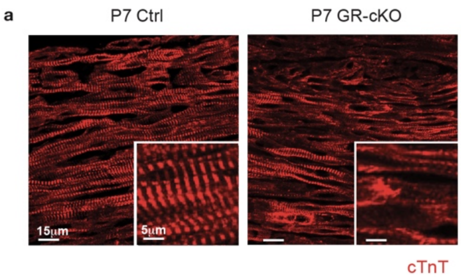


**Supplementary Figure 7. Sarcomere status of P7 GR-cKO and control cardiomyocytes.** *In vivo* cardiomyocyte sarcomeric status evaluation by immunofluorescence analysis of cardiac troponin T (cTnT) in P7 control (ctrl) and *GR*-cKO heart sections. Images were obtained using a confocal microscope.

**Supplementary Table 1.** Sequences of the primers used for mouse genotyping analyses.

| Gene | Forward primer | Reverse primer |
|---|---|---|
| αMyh6-Cre | ATGACAGACAGATCCCTCCTATCTCC | CTCATCACTCGTTGCATCATCGAC |
| αMyh6-Cre (internal control) | CAAATGTTGCTTGTCTGGTG | GTCAGTCGAGTGCACAGTTT |
| $GR^{flox}$ | ATGCCTGCTAGGCAAATGAT | TTCCAGGGCTATAGGAAGCA |

**Supplementary Table 2.** Sequences of the primers used in this study to analyse mRNA levels by real-time (rt)PCR.

| Gene | Forward primer | Reverse primer |
|---|---|---|
| NR3C1(GR) | TGATGTGGTTGAAAATCTCC | GTAAGGCAGTCATTTCTGATG |
| TNNT2(cTnT) | AGAGTATTCACAACCTGGAG | GAGTTTTGGAGACTTTCTGG |
| PECAM | CATCGCCACCTTAATAGTTG | CCAGAAACATCATCATAACCG |
| DDR2 | CGAGATCACTTTCCAATCAG | ACAGGATGATGACGATGATAG |
